# RealTime Heart Rate Monitoring Using Photoplethysmographic (PPG) Signals During Intensive Physical Exercises

**DOI:** 10.1101/092627

**Authors:** Majid Farhadi, Mahdi Boloursaz Mashhadi, Mahmoud Essalat, Farokh Marvasti

## Abstract

Heart Rate (HR) is a fundamental vital sign, monitoring which provides essential information for automated healthcare systems. The emerging technology of Photoplethysmograph (PPG) is shown as a feasible candidate for such applications; however, Motion Artifacts (MA) hinder efficient HR estimation using PPG, especially in situations involving physical activities. It is previously shown that even in the presence of sever MA, HR is still traceable with the help of simultaneous acceleration data although at high computational expenses. In this paper, we propose a novel framework, that not only improves the accuracy in HR estimation, but also achieves realtime performance by significantly reducing the complexity of system; mainly due to alleviation of the need for computationally demanding MA cancellation methods. Utilizing an spectrum estimation model (autoregressive) that suits well to the inherent PPG generation process, and benefiting from further intrinsic properties of the environment (e.g., the venous pulsation phenomenon); our framework achieves realtime and delayed (post-processed) Average Absolute Errors (AAE) of 1.19 and 0.99 Beats Per Minute (BPM) respectively, on the 12 benchmark recordings in which subjects run at speeds of up to 15 *km*/*h* maximum. Moreover, the system makes standalone implementation feasible by processing input frames (2 channel PPG and 3D ACC) in < 0.004 times of the frame duration, operating on a 3.2 GHz processor. This study provides wearable healthcare technologies with a robust framework for accurate HR monitoring; at considerably low computational costs.

## I. Introduction

Healthcare systems are of particular interest during physical activities especially when patients need continuous medical measurement of vital signs, among which, *Realtime* Heart Rate (HR) monitoring is a fundamental assess. In this paper we propose a framework to provide accurate and realtime HR monitoring, at significantly reduced computational cost to make it feasible for standalone operation on conventional wearable devices.

Using Electrocardiography (ECG) - detecting Heart Beat (HB) correlated electrical signals from the skin - has been a traditional method to accurately determine the HR. However, requirement of ground and reference sensors, vulnerability to nearby electric devices, portability issues, and implementation costs are ECG susceptibilities that pave the way for substitute technologies, e.g., Photoplethysmography (PPG) [1], [2]. Simply speaking, PPG measures the amount of light reflected from the skin. PPG signals are correlated with the alterations in blood pressure, and thus contain HR information [3]–[5].

Vulnerability of PPG to artifacts caused by body movements during the recording play as the Achilles heel for PPG. Many previous works [6]–[12] proposed signal processing algorithms for MA reduction in weak MA scenarios such as finger movement and walking. Moreover, some of them that address the problem of HR monitoring using PPG signals during intensive exercise scenarios, consider low MA PPG signals recorded from fingertip or ear.

An effective technique to cleans Motion Artifact (MA) contaminated PPG signals is to utilize simultaneously recorded accelerometer (ACC) signals as MA reference [13], [14]. In this paper we consider the problem of efficient HR estimation, during intensive physical exercises, using simultaneous wrist-type PPG and 3D ACC signals.

This line of research was initiated by the seminal works of Zhang et. al. [2], which proposed the TROIKA framework, consisting of three main steps: signal decomposition by Singular Spectrum Analysis (SSA), Sparse Signal Reconstruction (SSR), and Spectral Peak Tracking. In TROIKA, SSA is used for denoising and sparsifying PPG spectrum, SSR yields a high-resolution estimate of the spectrum, and spectral peak tracking step selects the peaks corresponding to HR by some decision mechanisms. This work achieved an AAE of 2.34 BPM on its proposed 12-dataset, which is also used as benchmark for performance comparisons by subsequent studies. Zhang [15] further improved this result by introducing the JOint Sparse Spectrum estimation (JOSS) technique. JOSS utilizes the Multiple Measurement Vector (MMV) model for simultaneous estimation of the spectra for PPG and Acceleration signals. Due to the common sparsity constraint on spectral coefficients, this method can identify and remove the peaks of MA in PPG spectra. JOSS achieved an AAE of 1.28 BPM on the benchmark 12-dataset. Although these works achieved acceptable performances in terms of the estimation error, the computational burden imposed by both SMV (Single Measurement Vector) and MMV sparse recovery algorithms make them less suitable for implementation on wearable devices.

In another research Sun et. al. in [16] proposed the SPEC-TRAP technique that removes the MA spectral components from the PPG spectrum using the asymmetric least squares technique. Subsequently, SPECTRAP uses Bayesian decision theory to locate the spectral peak corresponding to HR. This technique achieved an AAE of 1.5 BPM on the benchmark dataset. In comparison with JOSS, SPECTRAP reduced the computational complexity at the cost of increased estimation error. Zhu et. al. [17] proposed the MICROST technique which consists of three main steps of Acceleration Classification (AC), preprocessing, and heuristic tracking. MICROST further transacts the estimation accuracy with the algorithm runtime -resolving multi-second frames in order of several milliseconds– at the cost of increased average estimation error to 2.58 BPM.

Continuing this line of research Boloursaz et. al. in [18] proposed to use successive adaptive filter stages to remove MA from PPG signals. The reference MA signals used for adaptive filter stages where extracted by decomposing the 3D acceleration signals to independent components, using Singular Value Decomposition (SVD). This was observed to facilitate convergence of the adaptive filters and improved the estimation accuracy, by achieving an AAE of 1.25 BPM. Investigating the same problem [19]–[21] achieved AAE of 1.8, 1.77, and 1.96 BPM respectively.

Up to this point, references [22]–[25] held the record in accuracy of HR estimation, by accomplishing rather close AAE of 1.06, 1.05, 1.07, and 1.11 BPM respectively.

With several recent studies achieving feasible AAE in experiments; other criteria become significant in determining practicable methods. First of all, runtime plays a role for the algorithms to be implementable at feasible costs and performances. It can also be critical for patients to be monitored in realtime, especially during physical exercises. Nevertheless, computational burden of the proposed systems is unfortunately overlooked in most cases. As a matter of fact, there are a few prior works that report the simulation time [16]–[18], [22], among which our framework achieves superior accuracy and runtime, with the ability of *realtime* operation.

Another threat for the HR monitoring scenario in the literature is HR track loss. The risk is increased by expanded errors in consecutive frames. These systems need a mechanism to detect this situation; while due to their low MA initialization assumption [2], the burden of a halt in the middle of physical exercise will be inevitable for their users. Unlike some of the prior works [2], [15], [18], our proposed MA suppression step is independent from the previous frame estimations. Moreover, in contrast with all, our system does not seek for the HR in a narrow window around previous HR estimation. So the risk of expanded errors is magnificently reduced. To probe this in practice, we consider a challenging scenario -namely memoryless HR tracking– in which no information from other frames is used to resolve a frame, and achieve an acceptable AAE of 4.60 BPM. From a practical viewpoint, our algorithm can also be a proper candidate, to initialize an arbitrary HR tracker in the middle of physical exercise.

These improvements are due to utilizing 4 observations on the problem environment, as described in section II. The first observation justifies utilization of the parametric Auto Regressive (AR) spectrum estimation technique; that allows us to replace the computationally demanding sparse spectrum estimation techniques, proposed by a couple of prior researches [2], [15], with the rapid Levinson-Durbin algorithm. The second observation justifies the role of 3D ACC signals in MA detection that clears the way to innovate a fast, while effective, MA suppression technique; namely Spectral Division. Our denoising method benefits linear runtime (to input size) and transcends the previously proposed techniques, such as SSA [2], ANC [18] and asymmetric least squares [16] in runtime. The third observation is on existence and justification of HB second harmonic in the PPG signal, due to which we intend to replace the conventional PPG spectrum by the proposed CUMulative SPECtrum (CUMSPEC) to detect HR. CUMSPEC makes efficient use of the second HR harmonic to further amplify the spectral peak corresponding to HR. The final observation on the intrinsic properties of HR and MA spectral footprints, is considered in design of a robust decision mechanism for tracking the HR.

A final contribution of this work is composed of three techniques, devised to improve performance of any arbitrary HR tracking system. First, the justified quality increase of PPG spectrum by sparsifying it using the Iterative Method with Adaptive Thresholding (IMAT) [26]. Second, achieving a more accurate (though slightly delayed) estimation of HR by applying a small order median filter on the sequence of HR estimates. Last but not least, we propose a technique to improve performance of an HR tracking system, for particular applications and/or arbitrary criteria (e.g., a specific physical activity, a user, …), utilizing Genetic Algorithms (GA) on effective parametric constants of the system.

The rest of this paper is organized as follows. In Section II we state fundamental ideas in design of our framework, which is formally introduced in Section III. Section IV explains the three general techniques, devised for improving arbitrary HR trackers. In Section V we report performance of the proposed framework in simulations on available benchmark datasets. Finally, Section VI concludes this work.

## II. Observations

In this section we explain the key observations, that shape the fundamental ideas of the proposed framework.

### A. Spectral HB Peaks Smeared by MA

Due to periodicity of the target HB signal in short time-frames, we investigate the spectral properties of system inputs. A first observation on the spectrum of PPG signals under study is that the spectral peak corresponding to HR is smeared by MA, in severely contaminated frames. The simple periodogram, utilized for spectrum estimation by a couple of prior works, fails to resolve the HR peak in highly contaminated PPG frames. This may occur due to the fact that spectral peaks corresponding to HR are masked in the sidelobes caused by energy leakage from MA peaks. This phenomenon is due the assumption of zero values for samples outside of the observation frame, in periodogram [27]. Subsequently, we propose Autoregressive (AR) model to recover the spectrum,which is explained in Section III-A. Figure 1 depicts the spectrum estimations by periodogram and an Autoregressive (AR) model, for a typical PPG frame. Due to leakage effects of the simple periodogram, the HR peak is displaced by sidelobes of the dominant MA component. However, the AR technique has been able to accurately locate the HR peak.

**Fig.1.**
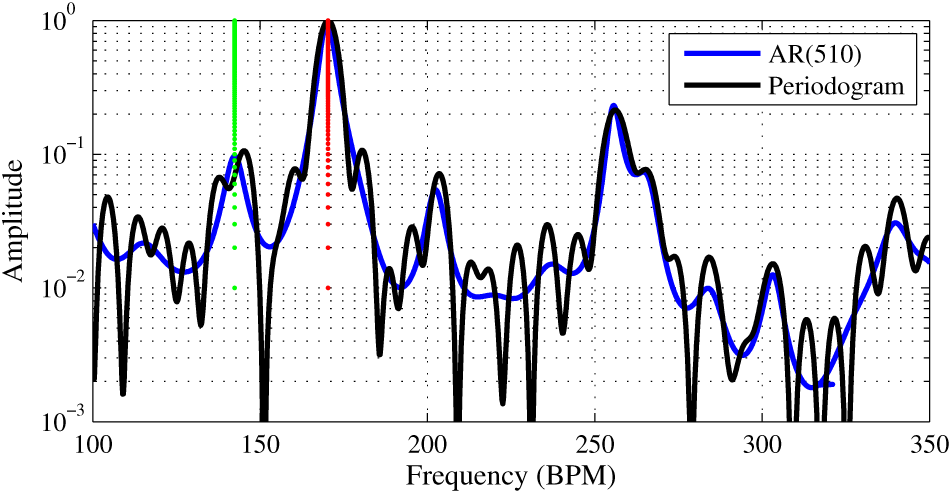
*HB Peaks Smeared by MA.* The spectral peak corresponding to HR (green) is displaced by a sidelobe of the MA peak (red) in periodogram. However, the AR technique deals well with this phenomenon.

### B. 3D-ACC Spectral Peaks as MA reference

The MA-contaminated PPG spectrum is affected by variations of the pressure between PPG sensor and the skin. In other words, the MA are caused by the periodic impulsive forces on the sensor-writs system, so it is highly correlated with 3D acceleration (ACC) data. In fact, the 3D-ACC signals include sparse spectral peaks that correspond to MA. These peaks are expected to be present in the PPG spectra and hence interfere with the HR peak as also observed in section II-A. Hence, a reasonable MA suppression approach is to extract ACC spectra from the PPG. To this end, we propose a fast MA Suppression technique named spectral division, which is further described in section III-B.

### C. Harmonically Related HR Peaks in PPG spectrum

A determinant factor on temporal shape of PPG signal, is the *venous pulsation* phenomenon [28], [29], creating a less strong but noticeable second peak in each cardiac cycle. As a result, HR is observed to possess a considerable second harmonic in the PPG spectrum as demonstrated in fig. 2, for data No. 2.

**Fig.2.**
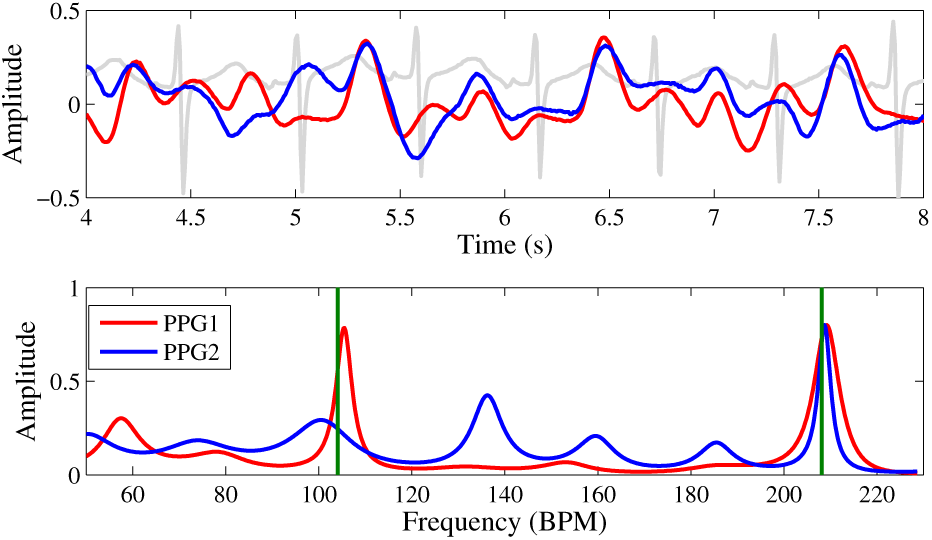
*Meaningful Harmonics*. Vertical lines mark HR 1^st^ and 2^nd^ harmonics.

Considering both harmonics in HR estimation, improves both accuracy and robustness in doing so. This approach can be redemptive especially for frames in which one of the harmonics is highly interfered by MA. To this end we propose an improved measure over the cleansed PPG spectrum, as stated in section III-C.

### D. Spectral Footprints of HR and MA

HB and MA frequencies pose different behaviors versus time. Figure 3 suggests this by over-sketching PPG spectra for several consecutive frames, in which HB-related peaks are gradually moving right while MA-related peaks show negligible mobility. Hence, the HR curve experiences a smooth and derivative-bounded plot. In contrast, it is observed that MA spectral peaks appear and vanish suddenly, but infrequently.

**Fig.3.**
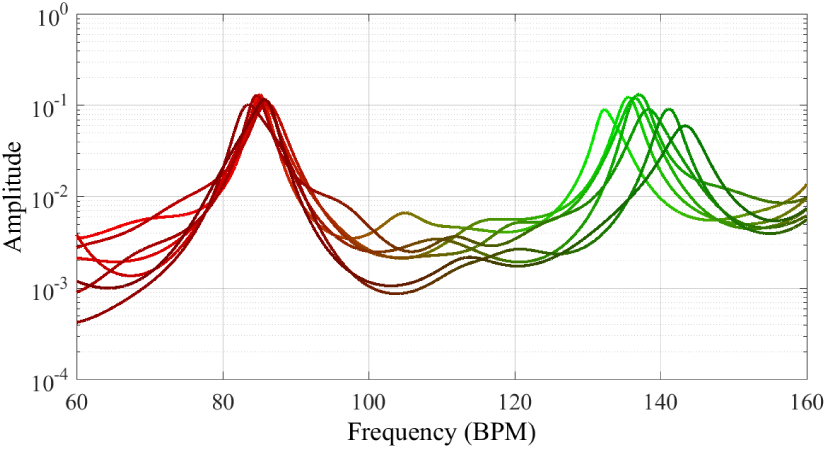
*Moving Peaks.* Spectra of *MA-contaminated PPG signals* in consecutive frames. Color code: HB correlated spectral peaks are colored in green, whereas the MA peaks in red.

Figure 4 also supports this claim by tracking time domain footprints of HR and the dominating MA frequency (maximizing sum of spectra for 3D-ACC), for another recording - No.12. To justify this claim, we note that the physical exercises usually consist of periodic body movements, each being performed for a limited duration. This observation shapes our attitude towards HR tracking during the exercise, expressed by the Lazy Tracker algorithm in section III-D.

**Fig.4.**
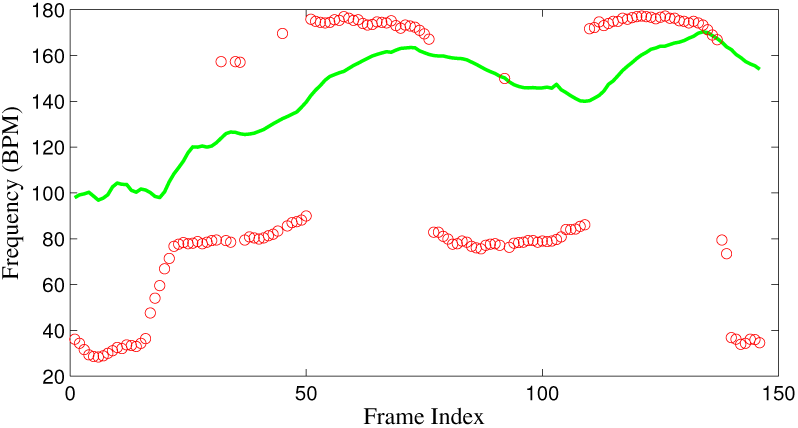
*HR vs MA*. Time plots for the ground truth HR (green), and dominating MA frequency (red).

## III. The Proposed Framework

Figure 5 presents the block diagram for our designed system, which is described in this section.

**Fig.5.**
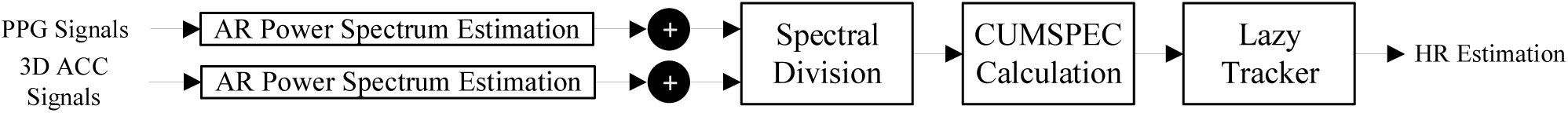
Overall Block Diagram of the Proposed Framework.

### A. AR Spectrum Estimation

Parametric spectrum estimation techniques can be used to minimize the previously mentioned side-lobe and leakage effects. Among different parametric techniques (AR, ARMA, MUSIC,…) the AR spectrum estimation technique is known to minimize sidelobes caused by windowing and provides the flattest spectrum by maximum entropy [27]. This technique is also computationally efficient as it can be realized recursively by the Levinson-Durbin algorithm [27]. Hence, we adopt it for spectrum estimation. Furthermore, in section V-E we prove superior performance of the AR technique in comparison with ARMA and MUSIC by simulations.

The AR model assumes input signals to be generated according to the linear difference equations in which *e*(*n*) is an input excitation sequence with a flat spectrum, *p*(*n*) denotes the PPG signal and *P* is the AR model order.

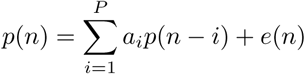

The estimated spectrum is an all-pole transfer function given by Equation (1), in which the unknown coefficients *a_i_*\* are iteratively calculated by the Levinson-Durbin algorithm as explained in [27].

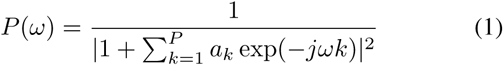

It should be noted, in the spectrum estimation step the algorithm estimates spectra for both PPG channels ({*P_i_*(*ω*)}) and the 3D ACC signals ({*A_i_*(*ω*)}). To make efficient use of diversity, we normalize the 5 estimated spectra and add them to achieve a single PPG and a single ACC spectrum, namely *P*(․) and *A*(․). Finally, note that normalization is required to cope with the effects caused by different PPG and ACC sensor gains.

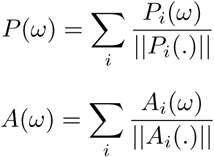

### B. MA Suppression by Spectral Devision

Continuing our discussion from section II-B, we intend to design an MA suppression procedure that extracts *A*(․) from *P*(․), with the aim to confront the situation in which MA energy is concentrated in sparse frequency peaks. Considering a threshold to accept the inevitable noise on ACC sensors, we intend to attenuate *P*(*ω*) in frequencies for which *A*(*ω*) is large. This step is roughly performed according to eq.(2).

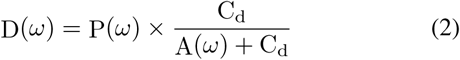

In eq.(2), D(*ω*) denotes the cleansed PPG spectrum. The constant C_d_, while helping to avoid division by zero, acts as a soft threshold to avoid attenuation by the small spectral ripples due to the inevitable noise on ACC sensors.

***Remark:*** The variance of sensor noise on the ACC signal spectrum can be an appropriate candidate for C_d_, yet we intend to train this parameter by optimization on a wide range, i.e., C_d_ ∈ (0,1).

### C. CUMSPECT measure calculation

While estimating HR by simply maximizing the amplitude in cleansed PPG spectrum is a common practice in the literature, this approach neglects the intrinsic second HB harmonic caused by venous pulsation that was mentioned in section II-C. However, in this subsection we propose the heuristic CUMSPEC measure that scores each frequency in the natural human HR range according to its own and second harmonic amplitudes in the cleansed PPG spectrum (eq. (3)) and propose maximization of this measure to determine the HR.

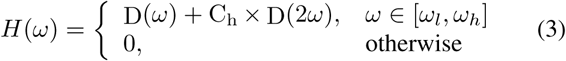

In eq. (3), H(*ω*) denotes the proposed CUMulative SPECtrum (CUMSPEC) measure, C_h_ is a constant weight and [*ω_l_,ω_h_*] covers human HR range.

***Remark:*** Any positive *C_h_* is expected to locally amplify *H*(*ω*) at ground truth HR (*ω_T_*) due to venous pulsation phenomenon. In contrast, a large *C_h_* can cultivate a misleading candidate peak at 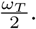 In this research we optimize *C_h_* in the range (0,2) as it is observed in low-MA PPG-frames that the ratio 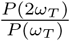 does not exceed 2. Although all *C_h_* in the specified range has shown promising performance in our experiments, its optimal value depends on specific constraints; among which the most influentials are arterial structure of the person and the part of body from which PPG is being recorded. Accordingly, *C_h_* increases the flexibility of our algorithm from two aspects. First, *C_h_* can be learned/adapted for a specific person, e.g., setting it to 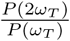 in low artifact frames. Second, it can help the algorithm to match a particular type of device, e.g., a headband or a wristband, because venous pulsation intensity varies for various parts of human body.

### D. HR evaluation by the Lazy Tracker

Considering the information within a frame `argmax`*ω_ω_H*(*ω*) will be an appropriate candidate for the system output, as has been highly correlated with ground truth HR in practice. Note that HR alterations are bounded during time and collision of HR and MA peaks in the spectrum is infrequent, so is the situation of sever HR attenuation in *D*(*ω*), where HR has no considerable peak in the cleansed PPG spectrum. To efficiently utilize this property we impose inertia on the peak tracking algorithm, namely the Lazy Tracker. This method searches for the dominating frequency in *H*(*ω*), in a wide spectral window where HR is expected to be in. Yet it performs a jump, bounded by a constant parameter (denoted by C_j_) towards that frequency. The Lazy Tracker is formally presented by pseudo-code in algorithm 1.

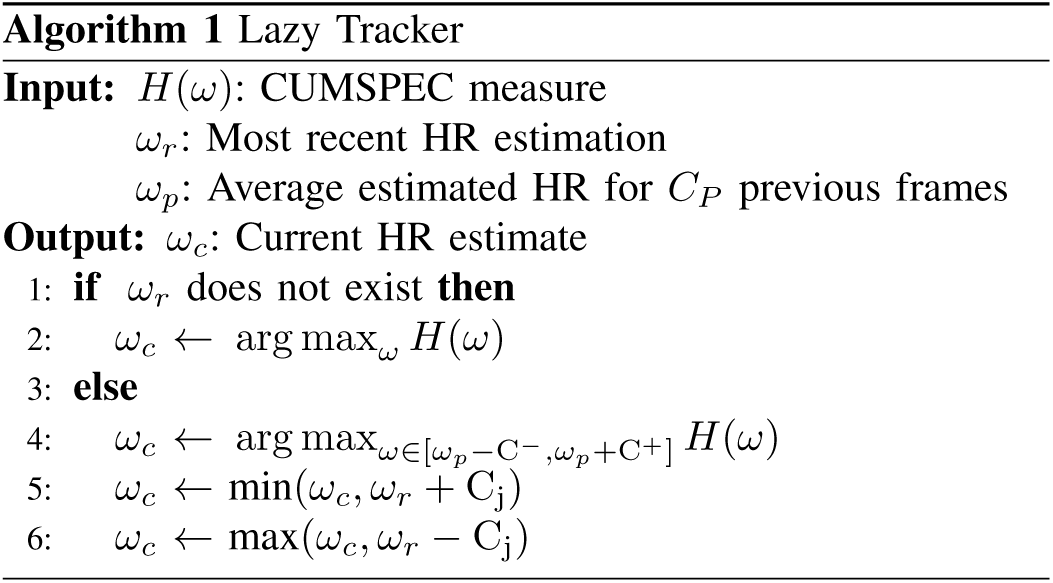

***Remark:*** Searching for HR in a wide window around *ω_p_* (average estimated HR for *C_*P*_* < 10 previous frames) improves the robustness of our algorithm, by enabling it to rescue back the track after severely MA contaminated frames. On the other hand, we need to limit HR variations between consecutive frames to avoid misleading jumps due to MA contamination. In order to meet both limitations we impose C^−^, C^+^ > 15 BPM and *C_j_* < 10 BPM.

## IV. Further Improvements

In the following subsections three general approaches are presented, that can improve performance of arbitrary HR tracking systems.

### A. Spectral Sparsification

Sparsity of cleansed PPG spectrum is a common and rational assumption in design of HR trackers. Sparsifying the cleansed PPG spectrum can enhance the position of spectral peaks. Considering our system as an example, although the maximum entropy AR model provides spectral estimates with minimal sidelobes, and the spectrum components corresponding to MA are attenuated by spectral division, there still exists some irrelevant spectral components that cause misleading scores to the subsequent CUMSPEC calculation step. In fact an ideal input to the CUMSPEC step in our algorithm is a sparse spectrum with sharp peaks at HR harmonics.

The task of further spectrum sparsification can be performed by the Iterative Method with Adaptive Thresholding (IMAT) algorithm [26]; given by eq. (4).

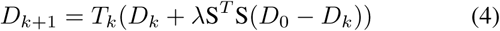

In Equation (4), *D*_0_ denotes the initial spectrum. *D_k_* is the sparsified spectrum estimated at *k*’th algorithm iteration. *S* is the partial Fourier matrix containing different harmonic exponentials. And *T_k_* is a thresholding operator decreased exponentially, i.e. *T_k_* = *T*_0_*e^−αk^*. The initial threshold value and its decay factor, *T*_0_ and *α*, were set to 0.7 and 0.87. We verified this idea by imposing 6 iterations of IMAT on the cleansed PPG spectrum (*D*(*ω*)) improving our realtime AAE on the 12 dataset to 1.17 BPM.

### B. Improved Delayed Estimation by Median Filtering

As described in section III-D, extreme scenarios such as exact collision of HR and MA spectral peaks can cause HB peaks attenuation in PPG spectrum, resulting outlier values in the estimated HR sequence. Some previous studies rationally faced this challenge by smoothening the sequence of HR estimates, e.g., moving average in [16]. To further improve the final report, we apply median filtering on the sequence of HR estimates to cancel outlier estimation values. Although this solution is non-causal, it is still practical if a delay equal to a few frames (two, for the simulation results reported below) in HR estimation is tolerable. Implementing this idea improved our AAE by 17%, down to 0.99 BPM, in the delayed (post-processed) estimation scenario as reported in table II.

### C. System Optimization

Performance of any system is affected by implementation details, e.g., its intuitively assigned constants. As well as previous studies, [2], [15], [17], [18], our algorithm contains effective parameters, e.g., C_d_ in spectral division, C_h_ in CUMSPEC calculation, and C^+^, C^−^, C_j_ in Lazy Tracker.

Instead of intuitive setup, we propose utilization of effective system parameters to attain optimal performance of the system. With more constraints on the environment this optimization becomes more powerful, e.g., HR monitoring during a special exercise, using a type of sensor, for a specific person.

This optimization problem can define an irregular and huge search space. So we propose to utilize artificial intelligence techniques for this challenge. Although using a simple hill climbing algorithm one can achieve a local optimum setting for a system, the realtime performance of our algorithm makes a diverse and powerful set of methods available, among which we chose evolutionary algorithms, particularly Genetic Algorithms (GA), due to their general assumptions on the search space. Modeling the vector of algorithm parameters as the gene and the resulting HR estimation error as the cost function, a GA inspects this multi-dimensional search space to minimize the cost function.

## V. Experimental Results

### A. Benchmark Datasets

To compare our framework with previous studies, we utilize a dataset that is widely used as benchmark in the literature. Provided by [2], the set includes 12 recordings from different healthful people, running on a treadmill for around five minutes at maximum speeds of 15 km/h. The signals (2×PPG, and 3D ACC) were recorded at 125 Hz from wrist. To enable a fare comparison with previous studies, HB is measured and reported in 8 *sec* time frames, each overlapping in 6 *sec* with the previous one.

We also report our performance on an additional 11-dataset provided for the second IEEE Signal Processing Competition, which contains a more challenging set of physical exercises. Details on the whole 12 + 11 dataset are represented in table I as reported by [2] and [23].

**TABLE I.** TEST SUBJECTS INFORMATION.

| Rec | Sub. ID | Activity | Sex | Age/Weight/Height | Health Cond. |
| --- | --- | --- | --- | --- | --- |
| 1 | 1 | WR | M | — | — |
| 2 | 2 | WR | M | — | — |
| 3 | 3 | WR | M | — | — |
| 4 | 4 | WR | M | — | — |
| 5 | 5 | WR | M | — | — |
| 6 | 6 | WR | M | — | — |
| 7 | 7 | WR | M | — | — |
| 8 | 8 | WR | M | — | — |
| 9 | 9 | WR | M | — | — |
| 10 | 10 | WR | M | — | — |
| 11 | 11 | WR | M | — | — |
| 12 | 12 | WR | M | — | — |
| 13 | 13 | RA | M | 20y 64kg 162cm | — |
| 14 | 14 | RA | M | 29y 70kg 169cm | — |
| 15 | 15 | RA | M | 21y 77kg 188cm | — |
| 16 | 15 | IA | M | 21y 77kg 188cm | — |
| 17 | 16 | IA | M | 19y 54kg 174cm | — |
| 18 | 13 | IA | M | 20y 64kg 162cm | — |
| 19 | 17 | IA | M | 20y 57kg 174cm | — |
| 20 | 18 | RA | M | 19y 70kg 180cm | — |
| 21 | 18 | IA | M | 19y 70kg 180cm | — |
| 22 | 19 | IA | M | 21y 73kg 180cm | — |
| 23 | 20 | RA | F | 58y 70kg 156cm | HR, BP |

Activities:
WR: Walking and Running on treadmill
RA: Rehabilitation Arm exercises
IA: Intensive Arm exercises, e.g. boxing.

### B. AAE Performance

Table II presents mean errors achieved by the proposed framework on benchmark datasets, for realtime and delayed estimation scenarios; outperforming previous studies. Last two columns of the table comprise our realtime and delayed (by utilizing a median filter of order 5 on the HR sequence) mean errors for the datasets.

**TABLE II.** AAE EVALUATION.

| Rec. # | [2] | [15] | [22] | [23] | RealTime | Delayed |
| --- | --- | --- | --- | --- | --- | --- |
| 1 | 2.29 | 1.33 | 1.83 | 1.23 | 1.81 | 1.51 |
| 2 | 2.19 | 1.75 | 0.85 | 1.26 | 1.44 | 1.25 |
| 3 | 2.00 | 1.47 | 0.63 | 0.72 | 0.63 | 0.55 |
| 4 | 2.15 | 1.48 | 1.21 | 0.98 | 1.16 | 1.01 |
| 5 | 2.01 | 0.69 | 0.65 | 0.75 | 0.83 | 0.74 |
| 6 | 2.76 | 1.32 | 1.03 | 0.91 | 1.40 | 1.28 |
| 7 | 1.67 | 0.71 | 0.70 | 0.67 | 1.02 | 0.88 |
| 8 | 1.93 | 0.56 | 0.50 | 0.91 | 0.63 | 0.50 |
| 9 | 1.86 | 0.49 | 0.47 | 0.54 | 0.68 | 0.57 |
| 10 | 4.70 | 3.81 | 2.83 | 2.61 | 2.77 | 2.16 |
| 11 | 1.72 | 0.78 | 1.14 | 0.94 | 1.03 | 0.78 |
| 12 | 2.84 | 1.04 | 0.90 | 0.98 | 0.90 | 0.68 |
| AAE-12 | 2.34 | 1.28 | 1.06 | 1.05 | **1.19** | **0.99** |
| 13 | — | — | — | 3.58 | 7.04 | 6.58 |
| 14 | — | — | — | 9.66 | 7.13 | 6.80 |
| 15 | — | — | — | 2.31 | 1.35 | 1.15 |
| 16 | — | — | — | 4.93 | 2.41 | 2.30 |
| 17 | — | — | — | 3.07 | 4.42 | 4.00 |
| 18 | — | — | — | 2.67 | 2.04 | 1.63 |
| 19 | — | — | — | 3.11 | 3.25 | 3.02 |
| 20 | — | — | — | 2.10 | 2.20 | 2.11 |
| 21 | — | — | — | 3.22 | 3.52 | 3.08 |
| 22 | — | — | — | 4.35 | 1.45 | 1.20 |
| 23 | — | — | — | 0.75 | 0.71 | 0.53 |
| AAE-23 | — | — | — | 2.28 | **2.17** | **1.93** |

In order to perform a fair comparison with previous studies, a fixed set of parameters is imposed on the whole dataset, i.e., (*C*_d_,*C_h_*,*C_p_*,C^−^, C^+^,*C_j_*)=(0.031,0.34,7, 25 BPM, 37 BPM, 5.1 BPM). These parameters are trained for minimized absolute error performance on data No. 10. This vector (modeled as a gene) is optimized by GA on a population of size 30, for 50 generations, while holding the constraints exposed in section III. Note that the training is performed on a single walking/running data and all other 22 tests, including various exercises and different persons, are *test data* for our experiment. This confirms robustness of the proposed optimization method while it still benefits flexibility for personalized performance, as will be described in section V-G.

***Remark:*** The basis for choosing data No. 10 to train our algorithm on was its challenging environment, e.g., compare the first twelve rows of table II. The ground truth HR curve for this recording experiences both steep and slow curvatures, while MA spectral peaks alternate around it. To verify this intuition we optimized the algorithm for every data in the 12 dataset and measured AAE-12 for the resulting settings. Trainings on tests 1,2,4,6,8,10 and 11 were successful, all with AAE-12 < 1.6 BPM, while the setting corresponding to data No. 10 holds the record. However optimized settings for the remaining five tests, that have been tracked more accurately than the other tests by previous studies, could not prevent failure on the whole dataset. This verifies the significance of training data quality.

### C. Runtime Performance

The complexity of our algorithm is asymptotically bounded by its spectrum estimation phase, while other framework stages benefit linear runtime complexities. MATLAB^™^ implementation for our system achieves realtime performance by processing 8 *s* frames in 28 *ms* (on average), running on a 3.2 *GHz* processor.

We intend to compare our runtime with the state-of-the-art algorithms mentioned in table II. Considering single-threaded MATLAB implementations, to normalize various processing powers, lets perform the comparison by Average Billion CPU Cycles Per Frame (ABCPF), being defined as follows.

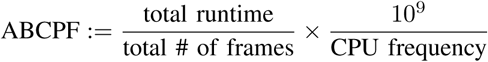

[2] and [22] perform in 25 and 1.36 ABCPF respectively, according to reports by [22], while our algorithm achieves runtime of 0.09 ABCPF. Considering the fact that modern smart watches are equipped with processors operating at clock rates higher than 0.5 GHz, the proposed algorithm seems suitable for stand-alone operation on wearable devices.

Although our framework achieves realtime performance, we can still trade between its speed and accuracy. Considering linear runtime of all phases in our system, except the spectrum estimation, we intend to perform runtime reduction by manipulating the resolution of reconstructed spectra. table III demonstrates this flexibility.

**TABLE III.** RUNTIME VS ACCURACY – THE 12-DATASET.

| Spectral Resolution | Runtime /Frame | Realtime AAE |
| --- | --- | --- |
| $2\pi/2^{15}$ | 28.1 ms | 1.19 BPM |
| $2\pi/2^{14}$ | 18.3 ms | 1.25 BPM |
| $2\pi/2^{13}$ | 13.7 ms | 1.29 BPM |
| $2\pi/2^{12}$ | 12.0 ms | 1.40 BPM |

### D. Modular Performance Investigation

We observe by experiments that a minimal subset of the designed system composed of MA Suppression and HR tracking stages is necessary and sufficient to avoid failure *(AAE > 20BPM).* Let’s call this the *Minimal* system, in which spectrum is estimated by simple periodogram and sparsity of PPG and HR second harmonic are disregarded. Table IV reports the AAE improvements when combinations of our proposed modules are utilized in the system.

**TABLE IV.** MODULAR PERFORMANCE INVESTIGATION — THE 12 DATASET.

| Algorithm | Realtime AAE |
| --- | --- |
| Minimal System | 1.82 |
| Minimal +CUMSPEC | 1.65 |
| Minimal +AR | 1.23 |
| Minimal +AR+CUMSPEC | 1.19 |
| Minimal+AR+IMAT+CUMSPEC | 1.18 |

### E. Investigating different spectrum estimation methods

We proposed to utilize the AR power spectrum estimation, due to observations expressed in section II-A. Similar discussions can expose other possible candidates for this phase, such as Auto Regressive Moving Average (ARMA (p,q)) [27] and MUSIC (p) [27]. Confirming our claims, ARMA and MUSIC proved superior compared to the simple periodogram by achieving AAE of 1.6 and 1.4 BPM, respectively. Finally, the proposed AR spectrum estimation technique shows superior performance among the three. It can be concluded that a periodic excitation HB signal filtered by the all-pole transfer function of the overall artery system and the PPG sensors best models the PPG signal generation process.

To benefit most from the AR model we intend to find the optimal AR model order. By considering different frame lengths (*n*∈{500,1000,2000}) we swept the AR order *p*. This experiment resulted a roughly concave AAE versus *p*, minimized at 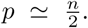 This is an expected result, because smaller *p* values tend to disregard less dominant spectral peaks which are highly probable to accommodate the HR. Also higher *p* values result in less accurate peak locations due to imprecise estimation of the lateral autocorrelation values.

### F. A Robust System

Memoryless functionality of the methods in our system -in spectrum estimation, denoising, and HR likeliness evaluation steps-reduces its vulnerability to distractions from true HR in previous frames. As an extreme case we propose the challenge of a complete memoryless HR estimation by performing initialization at every frame, in which our algorithm achieves an AAE of 4.60 BPM on the 12 dataset.

### G. Conditional Optimization

Result of performing the proposed GA optimization schema on individual records of the dataset is represented in Table V, where the manipulated parameters and their optimal values are also included. This experiment demonstrates flexibility of our framework to be specialized for various applications.

**TABLE V.** CONDITIONAL OPTIMIZATION.

| Rec. # | $C_h$ | $C_d$ | $C_j$ | RealTime | Delayed |
| --- | --- | --- | --- | --- | --- |
| 1 | .41 | $1.1 \times 10^{-2}$ | 3.0 | 1.34 | 1.14 |
| 2 | 1.3 | $3.1 \times 10^{-3}$ | 8.0 | 0.91 | 0.73 |
| 3 | 1.5 | $4.4 \times 10^{-2}$ | 5.1 | 0.58 | 0.55 |
| 4 | .34 | $4.0 \times 10^{-2}$ | 7.4 | 0.95 | 0.86 |
| 5 | 1.5 | $4.5 \times 10^{-2}$ | 7.0 | 0.76 | 0.71 |
| 6 | .53 | $1.6 \times 10^{-2}$ | 5.1 | 1.12 | 1.01 |
| 7 | .98 | $4.0 \times 10^{-1}$ | 3.2 | 0.70 | 0.59 |
| 8 | 1.2 | $1.6 \times 10^{-2}$ | 5.7 | 0.58 | 0.47 |
| 9 | 1.4 | $4.4 \times 10^{-2}$ | 3.2 | 0.58 | 0.50 |
| 10 | .34 | $3.1 \times 10^{-2}$ | 5.1 | 2.77 | 2.16 |
| 11 | 1.3 | $1.6 \times 10^{-2}$ | 6.3 | 0.91 | 0.70 |
| 12 | 1.6 | $3.0 \times 10^{-3}$ | 5.4 | 0.82 | 0.66 |
| Average Absolute Error |  |  |  | **1.00** | **0.84** |

## VI. Conclusion

In this paper we proposed a novel framework for realtime heart rate monitoring using simultaneous wrist-type PPG and 3D acceleration signals, when the subjects are performing intensive physical exercises. This framework consists of four main steps: AR spectrum estimation, MA suppression by spectral division, CUMSPEC calculation, and HR initialization/tracking using the proposed “Lazy Tracker” algorithm. We also devised general methods to further improve the performance of any arbitrary HR monitoring framework. In comparison with state-of-the-art algorithms in the field, our proposed framework shows admissible HR estimation accuracy in various test environments, including running and boxing exercises, while it benefits in considerably reduced computational costs and achieving realtime performance.

